# Measuring partner synchrony during salsa dancing and its relationship to changes in psychosocial domains

**DOI:** 10.1101/2025.08.26.672126

**Authors:** Molly Scott, Shelby Lam, Brittany Pereira, Grace Tsui, Mark Vrbensky, Filip Potempski, Aaron Wallace, Ilona Posner, Kara K Patterson

## Abstract

**Background:** Dance is a promising rehabilitation adjunct but understanding the mechanisms through which dance improves physical and psychosocial well-being is necessary for implementation. This study investigated the feasibility of a measure of one possible mechanism: partner synchrony. Secondary objectives were to examine the relationships between partner synchrony, instructor ratings, and changes in psychosocial variables.

**Methods:** Participants wore an Inertial Measurement Unit (IMU) sensor during a salsa class that was video recorded. Partner synchrony was quantified with correlations of x, y, and z-axis acceleration between dancing pairs. Psychosocial variables were measured pre- and post-class. Salsa instructors rated partner synchrony from video recordings. Feasibility parameters included percentage of participants with data collected and sensor comfort. Pre-post changes were analyzed with Wilcoxon ranked tests and relationships were analyzed with Spearman correlations.

**Results:** Data was collected for 23/24 (96%) participants and 17/24 (81%) reported the sensor was comfortable. All psychosocial variables improved from pre- to post-class. Partner synchrony was significantly associated with instructor ratings and change in positive and negative affect.

**Conclusions:** An IMU-based measure of dance partner synchrony is feasible, associated with ratings by dance instructors and related to changes in mood. The partner synchrony measure can be used to advance our understanding of dance as a therapeutic tool after its reliability is investigated.

## Introduction

Dance is a worldwide human activity that involves synchronized movement of the body to a rhythmic stimulus such as music (Quiroga et al., 2010). Dance has diverse effects in the physical and psychosocial domains across various populations (e.g., professional and recreational dancers, patients) (Chen et al., 2024). Physical effects of dance include more stable gait, better balance and aerobic fitness (Chen et al., 2024). Psychosocial effects include improved mood, reduced stress and social connection (Chen et al., 2024).

Given the numerous physical and psychosocial benefits, there is increasing interest in dance as a holistic rehabilitation adjunct for a variety of clinical populations including Parkinson’s disease and stroke (Kipnis et al., 2023; Patterson et al., 2018; Sharp & Hewitt, 2014). However, much work is needed to advance dance as a rehabilitation approach. In this context, dance fits the Medical Research Council definition of a complex intervention (Council, 2019). An important aspect of investigating and implementing a complex intervention is understanding the “active ingredients” or the physiological mechanisms through which it may have its impact (Council, 2019). The underlying mechanisms through which dance bestows its benefits have not been fully explored. It is possible that movement synchronization is one of likely many active ingredients of dance.

Movement synchronization with other people fosters feelings of social connection which in turn may reduce stress and improve mood. A wide array of synchronized physical activity in a group (e.g., chanting, moving limbs to a metronome cue, walking in step) has prosocial effects, including cooperative behaviour, trust, and entitativity (the degree to which a gathering of people are perceived as a group) (Reddish et al., 2013; Reddish et al., 2016; Wiltermuth & Heath, 2009). For example, groups of participants learned a series of movements and then performed those movements to music for 10 minutes either together in the same order (synchrony) or as a group with each person performing the movements in a different order (partial synchrony) (Tarr et al., 2015). The synchrony group exhibited greater increases in a composite prosociality measure that included dimensions of likeability, connectedness, and trust (Tarr et al., 2015). Furthermore, prosocial behaviour and positive affect reinforce each other in daily life (Snippe et al., 2018) and an interventional study revealed that prosocial behaviour, social connection, emotion, and physical health interact in a “self-sustaining upward-spiral dynamic” (Kok et al., 2013). Thus, inducing prosocial behavior and social connection through synchronized movement in dance may reinforce positive mood and overall health.

A deeper understanding of movement synchrony as an active ingredient will facilitate investigations into the effects and the implementation of dance-based rehabilitation interventions. For example, if we discover that improved mood and increased connectedness is related to synchronization with a dance partner (i.e., partner synchrony), this information could inform the design of dance interventions for patient populations such that partner work is emphasized to address social isolation and depression common with neurological conditions like stroke and Parkinson’s disease (He et al., 2025; Prenger et al., 2020; Rickards, 2005). Such investigations of the active ingredients of dance require a method to measure partner synchrony while dancing.

Wearable sensors have received significant attention and have demonstrated value in measuring movement in patient populations, including stroke (Gebruers et al., 2010) [16]. Furthermore, wireless accelerometers have been used to measure the movement of people dancing a variety of styles, including ballet (Thiel et al., 2014), jazz (Stančin & Tomažič, 2021) and Latin dance (Domene & Easton, 2014). However, the use of accelerometers in clinical settings presents challenges. For example, clinicians have highlighted the time to download and interpret data and a lack of protocols for device management as barriers to the implementation of accelerometer devices (Maher et al., 2021). In contrast, simple metrics were highlighted as enablers of the use of devices in clinical practice (Maher et al., 2021). Thus, to be useful for both clinical research and the clinical implementation of dance, a measure of partner synchrony will need to be simple and straightforward in interpretation.

The primary objective of this study was to determine the feasibility of measuring movement synchronization between dance partners (partner synchrony) during a single one-hour beginner salsa dance class with IMUs. Secondary objectives were to describe the relationships a) among measures of partner synchrony in the x, y, and z axes; b) between measures of partner synchrony and ratings of synchronization by salsa instructors; and c) measures of partner synchrony and changes in mood, perceived stress, and social connection after a salsa class.

## Methods

This study used a cross-sectional pre-post-test design. Ethics approval was granted from *blinded* Health Sciences Research Ethics Board.

### Participants

Neurotypical adults between 18 and 50 years of age were recruited from the surrounding community. Participants were included if they had no self-identified hearing deficits and no pre-existing neurological or orthopaedic conditions that significantly impacted mobility. Participants were excluded if they had pre-existing relationships with more than two members of the salsa class. All participants provided informed written consent.

### Experimental condition – beginner salsa dance class

Salsa (on-1 or LA-style) is a linear dance that involves a series of 6 steps taken over 8 counts in the music in either a front-and-back or a side-to-side pattern. Synchronization of the 6 steps to the beats of the music and with the dancing partner is emphasized in this style. This style also limits arm movement due to partnering, which requires hand holding. These characteristics of the Salsa on-1 style facilitate the measurement of gross body movement with accelerometry using a single wireless IMU sensor at the sternum.

The experimental condition was a single one-hour beginner salsa (on-1 style) class. Three classes in total were delivered on separate days to three groups of 8 participants. All classes were taught by the same Latin dance instructor (12 years of experience) using the same music and choreography. The format of the class, designed by the instructor with the knowledge of the study objectives. The class included instruction in the basic salsa steps, such as forward basic, side basic, and right turn, as well as basic partnering skills, including open hold and leader-follower skills for step changes and under-arm turns. Participants rotated partners to ensure that they were synchronizing their movements with multiple members of the class - a typical practice in salsa classes. These intervals of partner dancing were 1-minute in length to ensure adequate data collection. Each participant completed three 1-minute partnered dance intervals (with 3 different partners) during which they performed the forward and side basic steps without turns. A pilot dance class was conducted to test the class template and music playlist prior to data collection to ensure consistency between the three salsa classes. The pilot data was not used in the final analysis. Participants wore a number pinned to the back of their shirts, and the classes were video-recorded to confirm consistency between classes and for offline rating of movement synchronization between dancing pairs by salsa instructors.

### Measures of interest

a. Demographics: Demographic information including age, sex, previous dance experience, and pre-existing relationships with members of the class was collected with a questionnaire before the salsa class.
b. Feasibility: Feasibility of measuring movement synchrony between dance partners while dancing was defined *a priori* by domains considered important for using the method in future work on dance as a complex intervention. Specifically, the following variables were tracked

i. percentage of participants for whom data was collected at a sample rate of 1024Hz and could be analyzed
ii. percentage of participants that responded yes to the question “Was the sensor comfortable to wear while dancing?”
iii. number of adverse events during the class or pre- and post-testing (i.e. falls, injury).

Thresholds for feasibility were set a priori as 80% of participants for both data collection and comfort and 0 for adverse events.
c. Psychosocial measures: Psychosocial measures were administered before and after the salsa class as paper and pencil tests. Change scores for each measure were calculated as change score = post class score – pre class score.

i. Mood was measured using the Positive and Negative Affect Schedule (PANAS) which is a 20-item self-report measure with good reliability and validity in a non-clinical population (Crawford & Henry, 2004; Watson et al., 1988). The PANAS is also sensitive to change within a single dance class (West et al., 2004). Participants rated each test item based on the extent to which they experienced each mood state in the present moment. Separate scores for the Positive Affect (PAS) and Negative Affect (NAS) states are calculated separately and range from 10-50, with higher scores representing higher levels of positive/negative affect.
ii. Perceived stress was measured using the Perceived Stress Scale (PSS) which has high internal reliability, is sensitive to change within a single dance class and is recommended for research (Cohen et al., 1983; West et al., 2004). Test items are scored on a four-point ordinal scale indicating how often a participant felt or thought a certain way in the past month. Scores range from 0-40 and a higher score is related to greater perceived stress.
iii. Social connectedness was measured using the Inclusion of Community in Self (ICS), which is a single-item pictorial measure that quantifies the feeling of connectedness to a community or group (Mashek et al., 2007). Participants are presented with a series of six images that feature two circles, one representing the community (salsa class) and one representing themselves, arranged in increasing degrees of overlap. A numerical value from one to six was assigned to each of the images for scoring: one being the least perceived social connectedness depicted by the most distanced circles, and six being the most social connectedness depicted by fully overlapped circles. The ICS is valid and reliable and correlates well with other measures of sense of community (Mashek et al., 2007).
iv. Comfort was measured after the class by asking participants to respond yes or no to the question “Was the sensor comfortable to wear while dancing?”

### Movement tracking during the salsa class

Gross body movement of each participant was recorded during the class using wire-less Shimmer3 IMUs (Shimmer Sensing, Dublin, Ireland). The IMU was fastened over the sternum with a chest strap with care taken to ensure the IMUs were oriented the same way on all participants, with the vertical axis (y-axis) aligned with the long axis of the trunk while standing erect. The Shimmer3 IMU is 51mm x 34mm x14mm and weighs 23.6 grams (Sensing, 2022). It contains a tri-axial accelerometer and tri-axial gyroscope (not used in the present analysis). The default axis directions for the Shimmer3 are as follows: x-axis aligns with mediolateral movement, z-axis aligns with anterior posterior movement and the y-axis is vertical (up and down) (Sensing, 2016).

Given the exploratory nature of this study, the sampling rate of the Shimmer3 IMUs was set to 1024Hz. All 8 IMUs were linked to the same timestamp on the laptop operating the manufacturer software ConsensysPRO (Shimmer Sensing, Dublin Ireland). The IMUs were synchronized by grouping and moving them together in the X-axis on a desktop prior to attachment to participants (Kluge et al., 2017). The salsa classes were observed by the researchers to monitor data collection and to use the ConsensysPRO event marker tool to mark the 1-minute time intervals when partner dancing occurred. The synchronization activity with the IMUs was also marked with the event marker tool. These markers were used later during offline analysis.

### Salsa instructor rating of movement synchrony

Construct validity refers to “whether you can draw inferences about the measurement values related to the concept being measured” (Heale & Twycross, 2015). Assessment of validity is an ongoing process and a judgment of the degree to which various forms of evidence support the interpretation of a measure (Strauss & Smith, 2009)]. As a first step to establishing construct validity of the partner synchrony measures, ratings of movement synchrony for dancing pairs were obtained from salsa instructors for comparison. One-minute excerpts from the video recordings of each of the three dance classes were created. These excerpts featured the partnered dancing from each class and aligned with the 1-minute of accelerometer data extracted for the calculation of the partner synchrony measures. The videos were in the frontal plane (back of one partner and front of the other partner), and the whole body of all 4 dancing pairs were in view. Dance instructors with at least 3 years of experience teaching Latin dance styles were recruited to rate how well dance partners synchronized their movements with each other. Instructors completed an eligibility questionnaire, viewed one of the 3 videos (randomly assigned), and then submitted their ratings of partner synchrony for each pair on a scale from 0 (moving independently) to 100 (perfectly synchronized) via a web-based questionnaire (Qualtrics, Provo, USA).

### Data Analysis

A custom code in MATLAB 2017 (MathWorks, Natick, USA) was used to calculate measures of movement synchrony between dancing partners using data collected from IMUs during the 1-minute partner dancing segments. Accelerometer data were filtered using a second-order Butterworth low-pass filter with a cut-off frequency of 5 Hz. A baseline offset was removed from the X, Y and Z data (using the MATLAB remove baseline offset function) based on the mean acceleration that was computed for the first identified time window of the class, with gravity also considered for the Y-axis.

The event markers were used to identify the start and end times of the sensor synchronization activity and the 1-minute intervals of partner dancing. The datasets were aligned between dancing partners based on those times.

### Partner synchrony – correlations of accelerometer signals in x, y, and z axes

Acceleration in the y-axis was the primary variable of interest for quantifying the synchronization of front-and-back and side-to-side salsa movement patterns between dance partners, as the center of mass would necessarily rise and fall in this direction when steps are taken in time with the music’s beat. However, accelerations in the x and z axes were also analyzed, given the main study objective to examine the feasibility of measuring partner synchrony.

Accelerations in each of the 3 axes over the 1-minute interval of partner dancing were correlated between dancing pairs using the MATLAB Corr function. The resulting correlation coefficients for the x, y, and z accelerations were assigned to both members of the partnership as an index of how well they synchronized their movement with each other. This was done for every 1-minute partner dancing interval. Thus, participants were assigned three correlation coefficients for each of the x, y, and z axes, one for each of the 1-minute partner dancing intervals they completed in class. (This measure was used subsequently for correlational analysis with the instructor ratings of synchrony, objective 2b). Then, the three coefficients for each axis were averaged within a participant to produce a mean x, y, and z value representing their ability to synchronize their movement to all partners during the class. The measures of partner synchrony were labelled PsynchronyX, PsynchronyY and PsynchronyZ, respectively.

### Statistical Analysis

Statistical analyses were performed using SAS 9.4 (Cary, USA), and significance was set at 0.05. The 3 dance classes were investigated for differences in demographics and baseline values for ICS, PSS, PAS, and NAS using one-way ANOVAs (ordinal data were rank-transformed prior to the ANOVA). Pre-post changes in psychosocial domains associated with the salsa class were verified using Wilcoxon signed-rank tests on change scores for ICS, PSS, PAS, and NAS. Effect sizes were also calculated as r = Z/√n (Tomczak & Tomczak, 2014). For objective 1 (feasibility of measuring partner synchrony) descriptive statistics were used to summarize feasibility data. Secondary study objectives were addressed with Spearman correlations among the accelerometer-based partner synchrony measures (PsynchronyX, PsynchronyY, PsynchronyZ) (objective 2a), between the correlation coefficients for acceleration in the x, y, and z axis between dancing pars and the salsa instructor ratings of synchrony (objective 2b), and between the accelerometer-based partner synchrony measures and change scores for ICS, PSS, PAS, and NAS (objective 2c).

## Results

### Participants

Twenty-four participants completed the one-hour salsa class. Table 1 summarizes demographics and pre- and post-class scores for the psychosocial measures for the whole group and each class. PAS (W =129, p<0.0001, r=−0.75) and ICS (W=115.5, p<0.0001, r= −0.82) significantly increased, and NAS (W=−78.5, p=0.0071, r= −0.52) and PSS (W=−79, p=0.0031, r= −0.56) significantly decreased from pre- to post-salsa class for the whole group.

**Table 1.** Demographics and psychosocial measures. Values presented are means (sd) for age or medians, and 1^st^ and 3^rd^ quartiles for psychosocial variables.

|  | Whole group | Class 1 | Class 2 | Class 3 |
| --- | --- | --- | --- | --- |
| Age | 24 (2) | 24 (2) | 25 (2) | 25 (3) |
| Gender F/M | 15/9 | 5/3 | 6/2 | 4/4 |
| IMU comfort<br>(yes/no/missing) | 17/4/3 | 6/2/0 | 7/0/1 | 4/2/2 |
| Pre PAS | 31.5 [28.0; 38.5] | 29.5 [27.0; 32.5] | 34.5 [29.0; 40.5] | 32.0 [28.0; 40.5] |
| Post PAS | 36.0* [33.5; 41.0] | 36.5 [33.5; 41.5] | 37.0 [40.5; 31.0] | 35.5 [35.0; 41.0] |
| Pre NAS | 14.0 [11.5; 17.5] | 15.0 [13.0; 22.0] | 14.0 [11.5; 20.0] | 12.5 [11.0; 14.0] |
| Post NAS | 12.0* [10.0; 14.0] | 13.0 [10.5; 14.0] | 12.0 [10.0; 14.5] | 11.0 [10.0; 13.5] |
| Pre PSS | 17.0 [14.0; 21.5] | 45.5 [14.0; 19.0] | 21.5 [17.5; 24.0] | 15.5 [10.5; 19.0] |
| Post PSS | 15.0* [10.0; 20.0] | 11.5 [10.5; 18.0] | 19.5 [12.0; 23.5] | 15.0 [7.5; 18.5] |
| Pre ICS | 3.0 [2.0; 3.5] | 3.0 [2.0; 3.5] | 2.25 [2.0; 3.5] | 3.0 [2.0; 4.0] |
| Post ICS | 4.0* [4.0; 5.0] | 4.5 [4.0; 5.0] | 4.0 [3.5; 5.0] | 4.5 [3.5; 5.0] |

Abbreviations: PAS = positive affect scale, NAS = negative affect scale, PSS = perceived stress scale, ICS = inclusion of community in self. \* Significant difference between pre and post class score for whole group, all $p < 0.01$

### Objective 1: Feasibility of measuring movement synchronization

No adverse events were observed during the three salsa classes. The Shimmer3 IMU units collected data at the desired sampling rate (1024Hz) for twenty-three (96%) participants. See Figure 1 for an example of accelerometer data for three dancing pairs.

**Figure 1.**
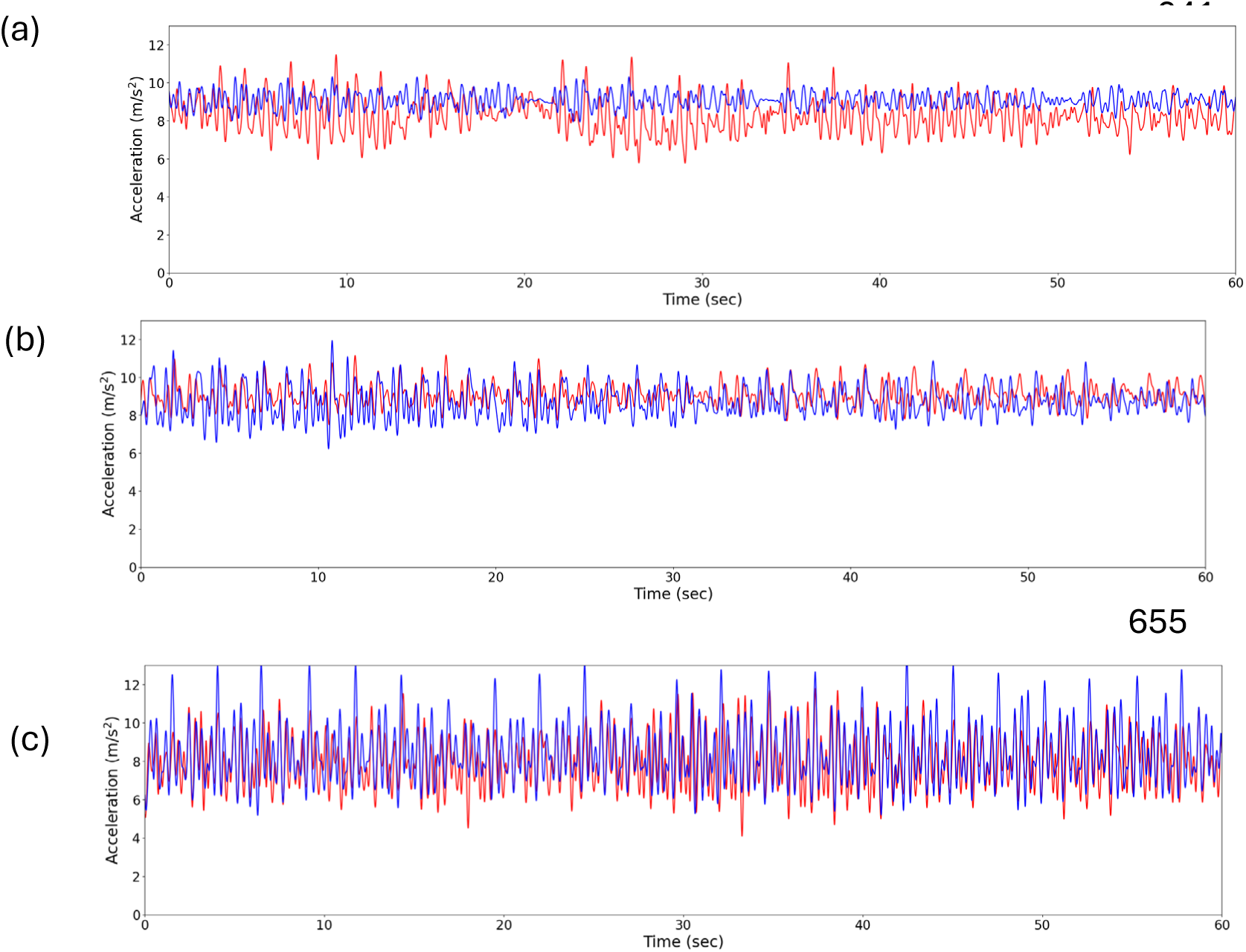
Accelerometer data in the y-axis for three dancing pairs (6 participants). (a) Dancing pair in class 1 with a partner synchrony of 0.42 and a mean (sd) instructor synchrony rating of 74 (20)%. (b) Dancing pair in class 1 with a partner synchrony of 0.001 and a mean instructor synchrony rating of 42 (18)%. (c) Dancing pair in class 3 with a partner synchrony of 0.28 and mean instructor synchrony rating of 65 (9)%.

An unknown error resulted in data collected at a lower sampling rate (51.2 Hz) for one participant. Minimal differences were observed when analysis was performed using a 1024 Hz sampling rate compared to a 51.2 Hz sampling rate. Therefore, the partnerships affected by the device with the 51.2 Hz sampling rate were down-sampled and analyzed the same way to produce partner synchrony measures.

Seventeen (81%) participants reported the Shimmer3 IMUs were comfortable to wear during the dance class (Table 1). Four (17%) participants reported the IMUs were not comfortable to wear; they were all men. A post hoc Fisher’s exact test was used to confirm this was a significant difference in IMU comfort between genders (F=0.0047, p=0.0085). Anecdotal comments made by participants who reported that the Shimmer3 IMU was not comfortable to wear while dancing revealed that discomfort arose from wearing the device over bulky or loose-fitting clothing such as button-down shirts or sweaters.

### Objective 2a: Relationships among accelerometer-based synchrony measures in 3-axes

The mean PsynchronyX, PsynchronyY, and Psynchrony Z values for the whole group and each class are summarized in Table 2. Only PsynchronyX and PsynchronyZ were significantly correlated (r_s_=0.63, p=0.001).

**Table 2.**
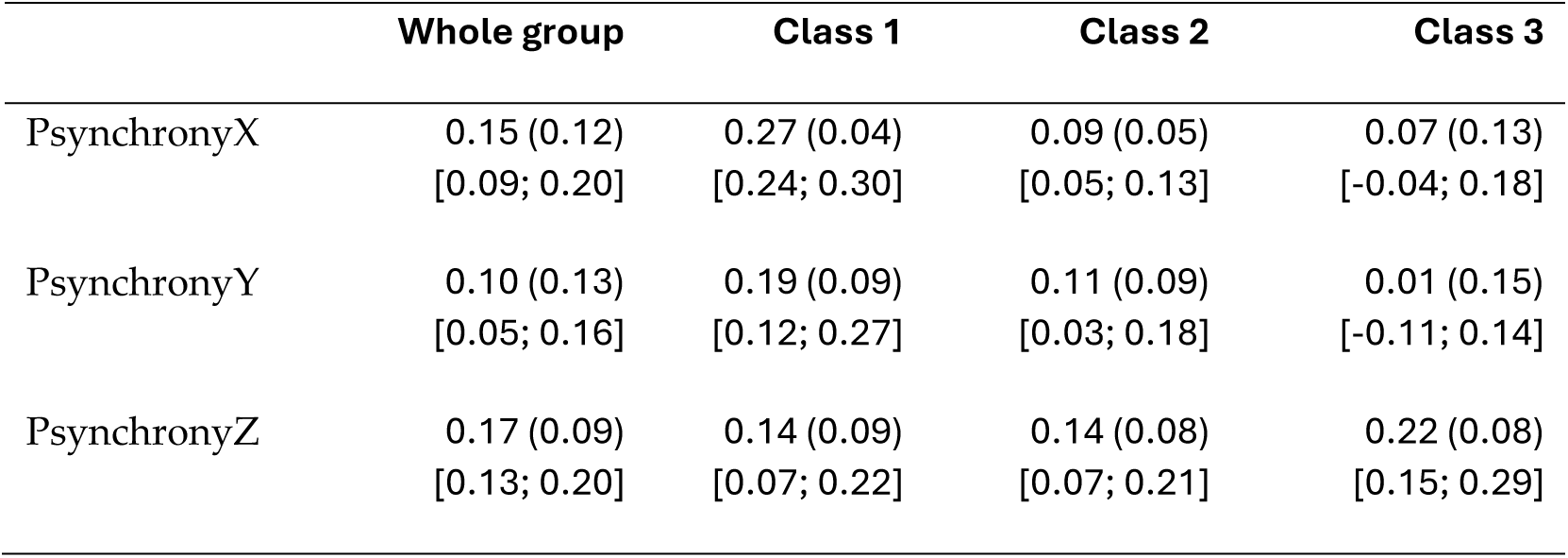
Accelerometer-based partner synchrony measures. Values displayed are mean (sd) and 95% confidence interval [CI] of the partner synchrony measures derived from correlations of the accelerations in the x (PsynchronyX), y (PsynchronyY) and z (PsynchronyZ) axes from each person in a dancing pair.

### Objective 2b: Relationship between accelerometer-based synchrony measures in 3-axes and salsa instructor ratings

Thirteen salsa instructors were recruited to rate the synchrony of dancing pairs from the video excerpts. Each dancing pair in the study was rated by 24 instructors. Instructor ratings of dance pair synchrony were significantly associated with the correlation coefficient of y-axis accelerations for the dancing pairs (r_s_ = 0.37, p = 0.0068). This relationship is illustrated in Figure 2. The salsa instructor rating was not significantly associated with the correlation coefficient for x (r_s_= 0.18, p= 0.19) or z (r_s_= 0.14, p= 0.3055) axis accelerations.

**Figure 2.**
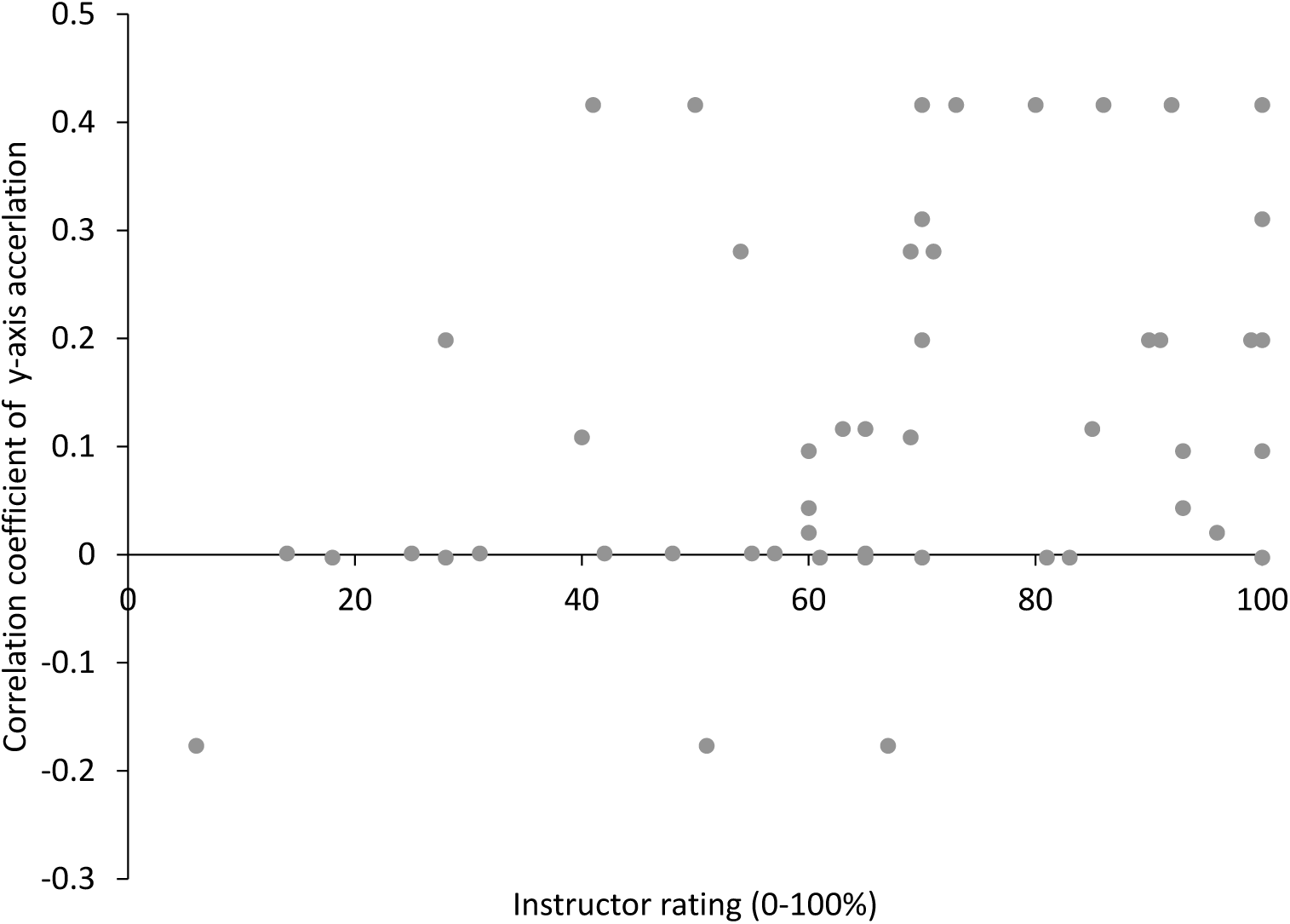
Relationship between partner synchrony in the y-axis and instructor rating of synchrony (r = 0.37, p = 0.0068). Each maker represents one dancing pair. Dancing pairs were rated by more than one salsa instructor who watched 1-minute video recordings of dancing pairs and rated synchrony on a scale of 0 (moving independently) to 100 (perfectly synchronized) via a web-based questionnaire.

### Objective 2c: Relationship between partner synchrony measures and changes in psychosocial measures

PsynchronyX was significantly associated with the PAS change score (r_s_= 0.41, p = 0.0494), and PsnchronyY was significantly associated with the NAS change score (r_s_= −0.41, p = 0.0455). These relationships are illustrated in Figure 3.

**Figure 3.**
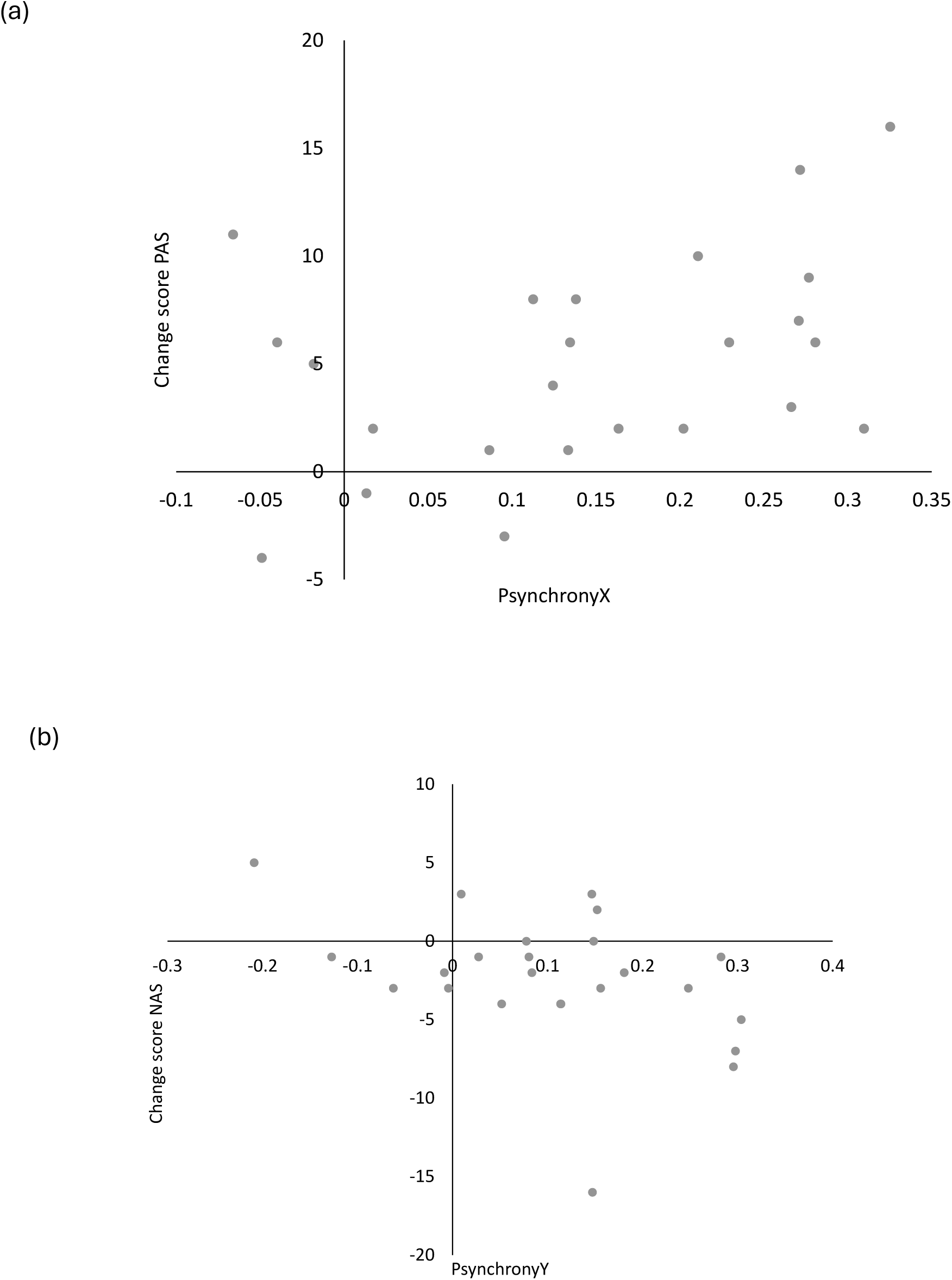
The relationship between PANAS changes scores and partner synchrony. Each marker represents one participant. (a) The relationship between PsynchronyX and change in PAS (rs= 0.41, p = 0.0494). Positive change scores represent and improvement or increased positive affect. (b) The relationship between PsynchronyY and change in NAS (rs= −0.41, p = 0.0455) with a salsa class. Negative change scores represent an improvement or decreased negative affect.

## Discussion

The main findings of this study are that it is feasible to measure partner synchrony with a single accelerometer during a salsa dance class and that change in positive and negative affect is associated with how well a person synchronizes their movements with their dance partners. To the best of our knowledge, this is the first study to measure partner synchrony during a dance class and characterize its relationship to changes in mood associated with that class.

The present results demonstrate the feasibility of measuring partner synchrony during a salsa class with a single IMU. Missing data is an important issue to consider with respect to the partner synchrony measure because eventually, it will be used in investigations of dance-based rehabilitation interventions. The reliability and interpretability of results from any clinical trial depend on the quality of data (Fleming, 2011). In the present study, data was collected for 96% of participants at the intended sampling rate. This aligns with the standards outlined in methodological assessment tools for clinical trials. For example, item 8 of the Physiotherapy Evidence Database (PEDro) scale specifies that measures of at least one key outcome are obtained for more than 85% of participants initially allocated to study groups (De Morton, 2009). In addition, item 3.1 of the revised Risk of Bias (RoB 2) tool notes that availability of data from 95% of participants is sufficient for continuous outcomes (Sterne et al., 2019). Due to an error, data was collected for one participant at a lower sampling rate (51.2Hz), requiring down-sampling of the data collected from this participant’s dance partners before the correlation analysis was completed. The lower sampling rate appears to have had little effect on the results, and thus, future work with the measure of partner synchrony can likely use a lower sampling rate. This aligns with previous work that used accelerometry to measure other features of movement (e.g., impact force estimates, tempo estimation, tilt of the torso) during other forms of dance (e.g., ballet, swing, jazz) at sampling rates from 100 to 500Hz (Almonroeder et al., 2020; Rossi et al., 2020; Stančin & Tomažič, 2021; Thiel et al., 2014). Finally, the present results suggest the setup and device (i.e., single IMU at the sternum) was comfortable for most participants, which is equally important to facilitate reliable and consistent data collection of partner synchrony. Anecdotal evidence from the current study also suggests that advising participants to wear thinner and more form-fitting clothing while dancing will improve comfort in future studies.

Synchrony between two people performing movements other than dance is commonly evaluated with correlations between two time series over short windows of interaction (Delaherche et al., 2012). In the present study this approach was applied to two series from the accelerometer signal recorded over 1-minute window of partner dancing. During salsa classes, dancers are instructed to synchronize their steps with the timing of the music which should also synchronize their steps with their dance partner if they are timing their movements to the music correctly. It is possible, particularly when the height is different between dance partners, for the timing of steps to be similar but the movement amplitude to differ. In this case, if the steps are occurring at the same time with respect to the music, the dancers would still be considered synchronized. Thus, correlation analysis is an appropriate method to analyze the similarity between accelerometer signals from a pair of dancers compared to other options such as coherence which provides an estimate of the overlap between movement frequencies in two signals (Guevara & Corsi-Cabrera, 1996). Furthermore, the present measure of partner synchrony in the y-axis correlated with ratings of synchrony by experienced salsa instructors. This is similar to previous work that used accelerometers to track the movement of ballet dancers performing a demi-plie to a counted rhythm and compared the accelerometer measures to the numerical scoring by a dance instructor (Thiel et al., 2014). This previous study reported Spearman correlation coefficients from r_s_=−0.64 to r_s_=0.63, which are larger in magnitude than the correlation reported in the present study (r_s_ = 0.37) but none the less, lends support to the construct validity of accelerometer-based measures of dance movement.

Further investigation is needed to test the reliability and validity of the partner synchrony measures. It will also be necessary to determine if analyzing accelerations in both the x and y axes are of value (given their different associations with the PAS and NAS measures) or if just the y-axis is sufficient given it was correlated with the synchrony measure in the x-axis. Regardless, this simple and straightforward methodology to directly measure partner synchrony during a group salsa class is promising as it facilitates investigating the active ingredients of dance as a holistic rehabilitation intervention.

As expected, there were improvements in mood, stress and social connection following the one-hour salsa class in the present study. Previous work has demonstrated that a single dance class or session reduces stress (West et al., 2004), improves mood (Quiroga et al., 2009; West et al., 2004), and increases social connectedness (Reddish et al., 2013). The present study extends these findings with direct, quantitative measures of partner synchrony, demonstrating that greater movement synchrony with dance partners in the mediolateral and vertical directions is associated with a greater increase in positive affect and a greater decrease in negative affect, respectively. Interestingly, partner synchrony was not associated with changes in stress or social connection. But the results of the present study do suggest that at least some of the psychosocial benefits of dance may be derived from the synchronization of movement with other people. In other words, partner synchrony may be one active ingredient of dance when implemented as a complex rehabilitation intervention. The potential effects of partner synchrony during a dance intervention on mood will need to be confirmed with a randomized controlled trial.

### Limitations

The small sample size may have limited the ability to detect significant relationships between partner synchrony and changes in the other psychosocial measures. Furthermore, participants were young, neurotypical adults and thus generalizability to older and/or patient populations is limited. Finally, although the construct validity of the partner synchrony measure was supported by an association with a subjective rating of synchrony by experts, accuracy and precision of the measures will need to be verified with an objective method such as video recognition or motion capture.

## Conclusions

This study demonstrates the feasibility of measuring partner synchrony during a salsa class with a single IMU sensor worn at the sternum. These results also add to our understanding of partner synchrony as a potential active ingredient of dance as a complex rehabilitation intervention that may reduce negative affect. Future work should determine the accuracy, reliability and validity of the partner synchrony measure which may be done through comparison to video recognition measures of footstep timing similar to previous work that matched footstep timing of a solo salsa dancer to the musical beat (Potempski et al., 2022).

## Author Contributions

All authors have reviewed and approved the submitted version of this manuscript. Author contributions are as follows: conceptualization AW, IP, KKP; methodology, AW, IP, KKP; investigation MS, SL, BP, GT, MV, AW; data curation, MS, SL, BP, GT, MV, AW, KKP; formal analysis MS, SL, BP, GT, MV, FP, KKP; project administration, AW, KKP; supervision AW, KKP, visualization MS, SL, BP, GT, MV; writing-original draft MS, SL, BP, GT, MV; writing-review & editing, MS, SL, BP, GT, MV, FP, AW, IP, KKP.

## Data Availability Statement

The datasets presented in this article are not readily available because ethics approval was not obtained for this type of data usage and/or dissemination

## Acknowledgments

This research was completed in partial fulfillment of the requirements for an MScPT degree at the blinded for review. The authors would like to acknowledge blinded for her time and effort creating and instructing the salsa dance classes.

